# Characterization of Onion Seedlots into Different Storability Groups with 4 Parameter Hill Function (4-PHF)

**DOI:** 10.1101/2020.08.17.253930

**Authors:** Sunil Kumar, Sudipta Basu, Anjali Anand, J. Aravind

**Affiliations:** ICAR-Indian Agricultural Research Institute, New Delhi -110 012; ICAR-National Bureau of Plant Genetic Resources, New Delhi -110 012

## Abstract

Onion varieties were classified into different storability groups by comparing two approaches (i)germination and vigor indices (conventional germination parameters) (ii)variables extracted from 4 Parameter Hill Function (4-PHF), after mimicking ageing conditions with accelerated ageing (42 °C and 100% RH). The study revealed that in comparison to evaluation using conventional germination parameters, the parameters extracted using 4-PHF provided realistic characterization of varieties as good, medium and poor storers. Time related parameters like time to maximum germination rate (*TMGR*), time to 50% germination (*T*_*50*_), difference between time at germination onset (*lag*) and 50% germination (*D*_*lag-50*_), uniformity (*U*) along with germination percent (*a*) and area under curve (*AUC*) were decisive in identification of the varieties to a storage category which was misinterpreted with exclusive use of conventional approach. The distinction between good and medium storers was not of much significance but shift of varieties like Bhima Super, Pusa Red and Agrifound Light Red from poor to good performance cluster could be detected exclusively through 4-PHF analysis. Curve fittings highlighted *AUC* as the most crucial parameter contributing towards clustering of the varieties in different storability groups. Our study is the first reported research of using 4-PHF mathematical function for seedlot characterization into different storability groups.

## Introduction

Seed longevity is described as the total duration of time upto which seeds stay viable or alive. It is an important trait, both agriculturally and ecologically, as it meets the food security and is required for regeneration of vegetation, respectively^1^.Seed deterioration bears an inverse relationship with seed longevity and is associated with gradual loss of quality (vigor, viability) and performance over time under storage conditions^2^. This leads to seeds becoming vulnerable to stress during germination and eventually death of seedling.

A considerable variability exists among the orthodox seed species w.r.t. seed longevity which is a complex and quantitative trait^2^.The rate of deterioration depends on storage conditions viz., temperature, relative humidity and oxygen pressure of storage units^3,4,5^. It is also affected by seed characteristics like seed moisture content, seed health and quality. Assessment of seed longevity under relatively dry and mild temperatures is a tedious process due to slow rate of deterioration. In order to hasten the process of deterioration, seedlots are placed under artificial ageing conditions i.e. high relative humidity and temperature. Artificial ageing treatments (controlled deterioration and accelerated ageing) mimic the natural ageing process and are widely accepted for studies related to seed longevity^2,6^. Seed sensitivity to artificial ageing tests has been successfully used to rapidly assess and predict seed vigor and longevity^6,7^.

Seeds of vegetable crops are expensive and loss of viability during poor storage conditions can account for huge economic losses for the seed companies and farmers. Among the vegetable crops, seeds of Allium sps., leek, lettuce, pepper and parsnip etc. are predisposed to losing viability if storage conditions are not controlled. Onion accounts for world’s second largest vegetable crop after potato with an area of approximately 5.4 million hectare and 103 million tonnes production. India is the second largest producer of onion after China and covers 16.93% and 19.25% of world’s area and production, respectively^8^. In India onion is grown in an area of 1.3 mha with a production of 23.28 million tonnes^9^.

The faster deterioration of onion seeds during storage is attributed to its fragile seed coat (cracks at hilum end) and chemical composition (lipid content; 22-26%)^10, 11^. The presence of high lipid content leads to increased probability of lipid peroxidation by the reactive oxygen species that accumulate during unfavourable conditions^12^.Characterisation of onion varieties as “good” and “poor” storers is essential to predict its deterioration during storage and recommend optimum packaging and storage conditions for improved longevity.

In most of the storage studies, only germination count is considered for assessment of seedlot quality, but seed germination is a dynamic process, the its outcome is determined by time, rate, synchrony and homogeneity^13, 14, 15, 16^. The speed or time spread of germination convincingly calculates the germination indices but misses on the “high or low germination events”^17^ that influences the vigor and stress resistance of the seedlots^18^.

Mathematical models have been used for characterizing the planting value of seedlots^15, 19, 20^ and 4-PHF is one such mathematical expression which uses multiple germination associated parameters to statistically interpret differences in germination capacity among seedlots^21^. The present study was conducted for classification of onion varieties into different storability groups by using the conventional approach (using germination count and vigor indices) *vis a vis* comprehensive analytical analysis by four parameter hill function(4-PHF) test. Our study is the first report on application of mathematical function i.e. four parameter hill function for classification of seedlots into different storability groups.

## Results

### Conventional approach using germination and vigor indices

#### Germination capacity (a)

Seedlots of seventeen onion varieties subjected to accelerated ageing test showed significantly different germination performance at distinct periods of ageing (48h, 96h and 144h). The germination percentage ranged from 91-100% in seedlots before ageing treatment, except in Agrifound Dark Red (87%). After 48h of ageing, the germination percentage of all varieties decreased, which further declined significantly after 96 and 144h of ageing ranging from 67-87% and 55-76%, respectively. Although genotypic differences persisted among varieties, Arka Niketan was an outlier at 96 and 144h ageing treatment exhibiting poor performance (germination percent of 59 and 40% respectively) (Fig.1a).

**Fig 1:**
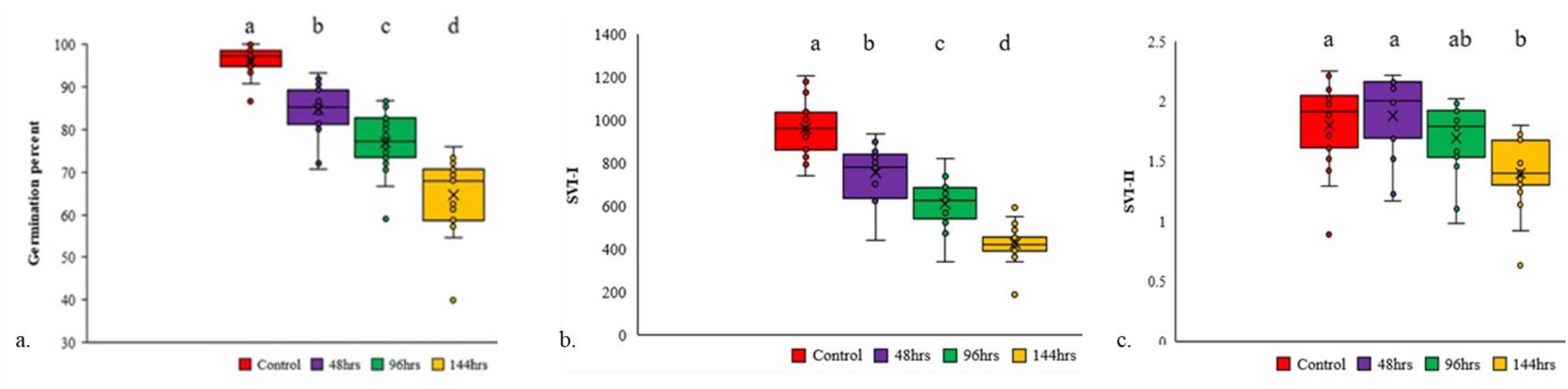
Boxplot showing the response of conventional germination parameters i.e.germination percent and seed vigour indices in unaged and 144 h aged seedlots at p≤0.05.

#### Seed Vigour Index-I (SVI-I)

The seedlots of seventeen onion varieties exposed to various ageing treatments exhibited significantly different seedling vigour index. SVI-I of unaged seedlots ranged from 743.06 - 1204.17 but progression of ageing to 46 and 96 h led to substantial decrease ranging from 442.82 - 902.06 and 342.12 - 821.05, respectively. Genotypic difference for SVI-I was clearly evident for all varieties at 144 h of ageing. Bhima Shwetha (1298.02) and Arka Niketan (190.62) showed the maximum and minimum SVI-I values (Fig. 1b).

#### Seed Vigour Index II(SVI-II)

Unaged and 48 h aged seeds did not show any change in seed vigour based on dry weight (SVI-II). Prolonged ageing upto 96 and 144 h caused a significant decline in the seed vigour 0.98-2.02 and 0.92 - 1.98, respectively. NHRDF Red (0.88) and Arka Niketan (0.63) were the poor performers at 48 and 144 h (Fig. 1c).

### Cluster analysis of the varieties based on germination percentage and vigor indices (SVI and SVII)

In order to check the stability in germination performance of different seedlots under unaged and aged conditions cluster analysis was done that helped in segregating the seedlots into 3 clusters. Here, heatmaps and dendrograms of the seedlots under unaged and 144 h of ageing are represented for examining the changes when the germination percentage was approximately reduced to 50% in most of the varieties. The characteristics of clusters under the two conditions are mentioned below:

Clustering based on the germination percent and seed vigor indices in unaged seedlots (fresh) (Fig 2: A.I & B. I)

i. Cluster 1 (Unaged) - It comprised of Bhima Kiran, Bhima Shakti, Bhima Shwetha, Bhima Shubhra, NHRDF Red varieties (Fig. 2: A.I&B. I). The mean values of germination percent and seed vigor indices showed that all varieties had *a* (97±1.4%), *SVI-I* (1130.75±34.78), *SVI-II* (1.63±0.20) as compared to other clusters (Table1).This cluster was designated as “good storer” as all the seedlots possessed high SVI-I
ii. Cluster 2 (Unaged) - Pusa Red, Pusa Madhavi, Pusa Riddhi, Agrifound Light Red, Arka Kalyan, Arka Bheem, Arka Ujjwal, Arka Pragati and Arka Niketan were placed under Cluster 2 (Fig. 2: A.I & B. I). Comparatively medium values of *a* (96±0.61), *SVI-I* (874.88±28.58) and *SVI-II* (1.82± 0.11) (Table1) were observed, therefore varieties were classified as “medium storers”.
iii. Cluster 3 (Unaged) - ADR, Pusa Early Grano, Bhima Super varieties were allocated to Cluster 3(Fig. 2: A.I &B. I). This cluster showed significantly lower *a* (90±3.33), *SVI-I* (905.16±56.55) and *SVI-II* (1.8±0.28) (Table1) that led to its classification as “poor storers”.

**Fig 2:**
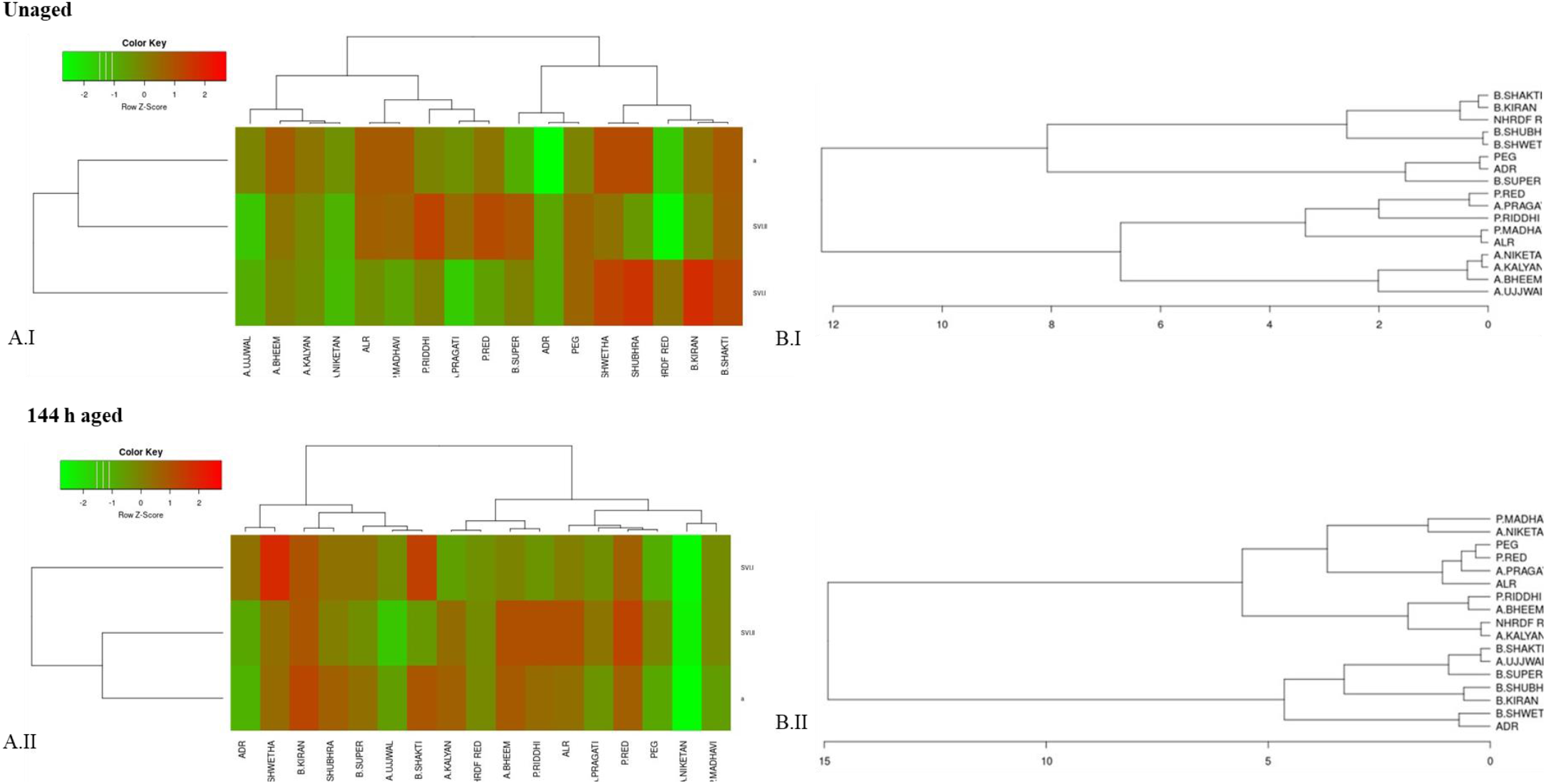
Heatmap (A) and dendrogram (B) generated from germination percent and vigor indices for clustering of varieties of (I) Unaged (II) 144 h aged seedlots. The data was subsetted per treatment with columns representing the value of individual samples and rows the selected parameter (a, SV-I and SV-II). Red and green represent high and low parameter value, respectively. The values of individual samples are normalized per parameter using z-Fisher transformation.

**Table 1:** Mean values of percent germination, seed vigor indices (I and II) used for classification of 17 seedlots of onion in different storability clusters (n=3). Cluster validation was performed by Tukey’s HSD pairwise test at p ≤0.05. Letters in superscript indicate significantly different groups

|  | **Unaged** |  |  |  |
| --- | --- | --- | --- | --- |
|  | **Varieties** | ***a* (%)** | ***SVI-I*** | ***SVI-II*** |
| **Cluster 1**<br>**(Good)** | Bhima Kiran, Bhima Shakti, Bhima Shwetha, Bhima Shubhra, NHRDF Red | 97±1.4<sup>a</sup> | 1130.75±34.78<sup>a</sup> | 1.63±0.20<sup>a</sup> |
| **Cluster 2**<br>**(Medium)** | Pusa Red, Pusa Madhavi, Pusa Riddhi, Agrifound Light Red, Arka Kalyan, Arka Bheem, Arka Ujjwal, Arka Pragati and Arka Niketan | 96±0.61<sup>a</sup> | 874.88±28.58<sup>b</sup> | 1.82±0.11<sup>a</sup> |
| **Cluster 3**<br>**(Poor)** | ADR, Pusa Early Grano, Bhima Super | 90±3.33<sup>a</sup> | 905.16±56.55<sup>b</sup> | 1.8±0.28<sup>a</sup> |
|  | **144 h aged** |  |  |  |
| **Cluster 1**<br>**(Good)** | Bhima Kiran, Bhima Shakti, Bhima Shwetha, Bhima Super, Bhima Shubhra, Arka Ujjwal and Agrifound Dark Red | 67±2.93<sup>a</sup> | 491.55±24.58<sup>a</sup> | 1.31±0.09<sup>a</sup> |
| **Cluster 2**<br>**(Medium)** | NHRDF Red, Arka Bheem, Arka Kalyan and Pusa Riddhi | 68±2.07<sup>a</sup> | 392.8±11.64<sup>b</sup> | 1.57±0.09<sup>a</sup> |
| **Cluster 3**<br>**(Poor)** | Pusa Red, Agrifound Light Red, Pusa Early Grano, Arka Pragati, Arka Niketan and Pusa Madhavi | 59±4.41<sup>a</sup> | 376.11±42.11<sup>ab</sup> | 1.38±0.16<sup>a</sup> |

Ageing for 144h resulted in classification of seedlots with differential storability in three different clusters (Fig 2 A. II & B. II)

i. Cluster 1(144 h aged)-It consisted of Bhima Kiran, Bhima Shakti, Bhima Shwetha, Bhima Super, Bhima Shubhra, Arka Ujjwal and Agrifound Dark Red (Fig. 2: A. II & B. II). The mean values of germination percent and seed vigor indices showed that all varieties had the highest *a* (67±2.93), *SVI-I* (491.55±24.58) and *SVI-II* (1.31±0.09) (Table1). This cluster was designated as “good storer” as all the seedlots possessed high viability/vigor.
ii. Cluster 2-It consisted of NHRDF Red, Arka Bheem, Arka Kalyan and Pusa Riddhi (Fig. 2: A. II & B. II). Comparatively medium values of *a* (68±2.07%), *SVI-I* (392.8±11.64) and *SVI-II* (1.57±0.09) (Table1) were observed hence, it was classified as “medium storer”.
iii. Cluster 3-It included Pusa Red, Agrifound Light Red, Pusa Early Grano, Arka Pragati, Arka Niketan and Pusa Madhavi (Fig 2 A. II&B. II). The cluster showed significantly lower *a* (59±4.41), *SVI-I* (376.11±42.11) and *SVI-II* (1.38±0.16) (Table1), therefore varieties belonged to “poor storer” category.

Cluster validation showed that *a (%)* and *SVI-II* were not significantly different amongst the different clusters under unaged and aged conditions. However, *SVI-I* only contributed towards classification of seedlots into good (cluster 1) medium (cluster 2) and poor (cluster 3) storers.

### Parameters extracted from 4PHF

The results of germination capacity (a) are presented in previous approach. In addition to that, the other extracted parameters were:

**Shape and steepness of the germination curve (b)** There was no significant difference amongst unaged and ageing treatments for shape and steepness of the germination curve (b). The *b* value in unaged (fresh seedlots) and 48h aged seedlots ranged from 2.58 −12.96 and 3.11 −11.98 respectively. After 96h of ageing it was 5.17 to 12.02 with one outlier Arka Pragati (18.87). After 144h of ageing, the range was 4.33 −8.99 with two outliers; Pusa Madhavi (9.76) and Bheema Shubhra (12.12) (Fig.3a).
**Time to 50% germination (T_50_)** There was significant difference between the unaged (1.73 to 5.26 days) and 48 h (2.23 to 3.44 days) aged with one outlier Arka Niketan (4.30 days). Whereas, the difference was non-significant after 96(2.49 to 4.05 days) and 144h (2.93 to 4.49 days) of ageing except in Arka Niketan (5.34 to 9.05days respectively) (Fig.3b).
**Difference between time at germination onset (lag) and 50% germination (D_lag-50_)** No significant difference was observed amongst the unaged and ageing treatments for the parameter being studied. D_lag-50_ in unaged and 48h aged seedlots ranged from 1.66 to 5.23days and 2.03 to 3.63days, respectively. Upon 96 and 144h of ageing, it spanned between 2.33 to 3.51 and 2.61 to 3.46 days respectively (Fig.3c).
**Time to Maximum Germination Rate (TMGR)** TMGR did not show a significant difference amongst the unaged and ageing treatments. It ranged from 1.46-5.12, 1.88-3.37 and 0.95-3.56days in unaged, 48 and 96h aged seedlots, respectively. Upon extending ageing to 144h, the TMGR shifted between 2.55 to 3.14 days (Fig.3d).
**Uniformity (U)** There was significant difference between unaged (1.08 to 5.26 days) and 48 h (0.9 to 3.6 days) of ageing treatment, but the difference was non-significant between the 96 h (0.95 to 3.56 days) and 144 h (1.01 to 5.13 days) of seed ageing (Fig. 3e).
**Area Under Curve (AUC)** The seed ageing treatments showed a significant effect on the area under curve in seventeen onion seedlots. Unaged and 48 h ageing treatments had no any significant difference in *AUC* of onion seedlots. Advancing the duration of ageing to 96 and 144 h led to the significant difference among the performance of varieties. The AUC of varieties ranged from 1196.860 - 1515.663 and 995.855-1298.017, respectively. Arka Niketan did not fall in the range noted for other varieties (Fig. 3f).

**Fig 3:**
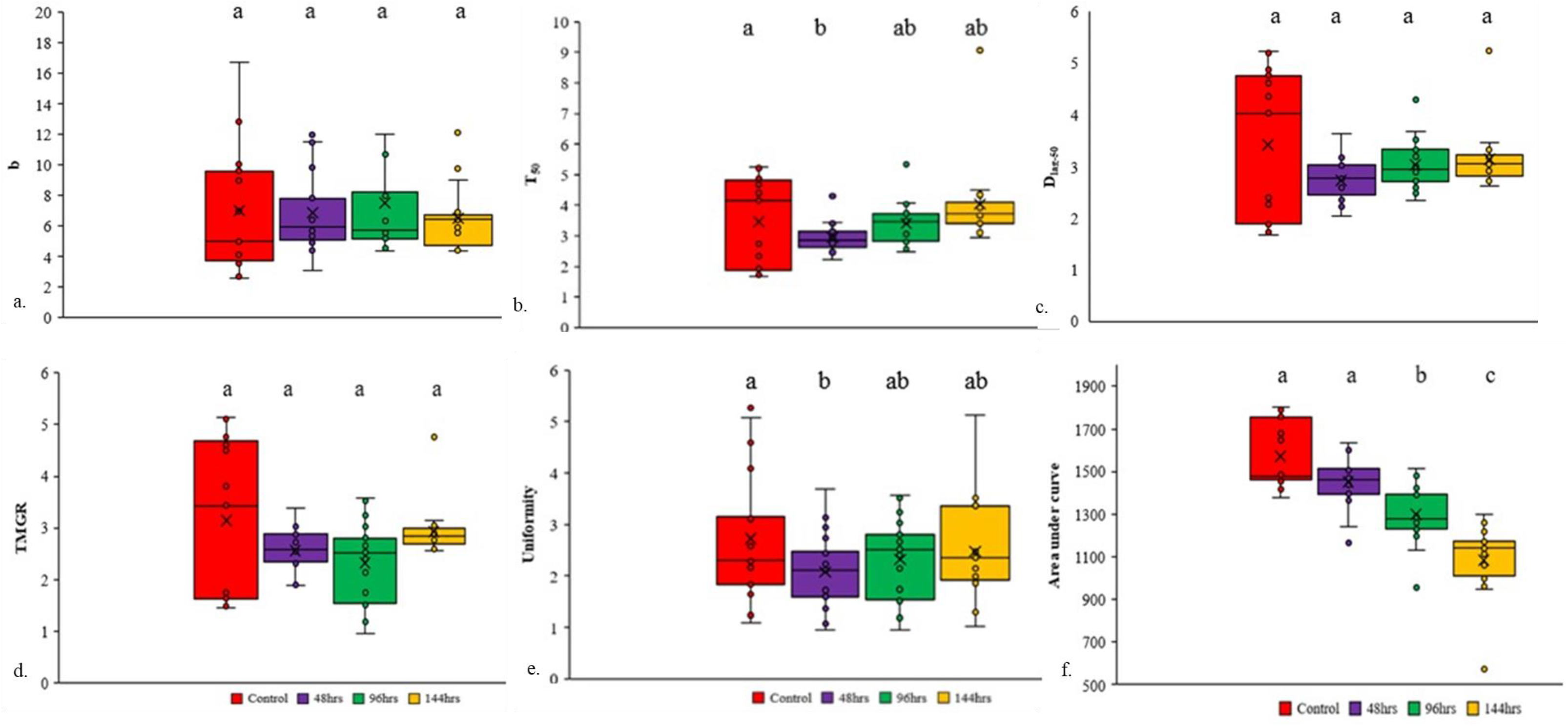
Boxplot showing the response of 4-PHF parameters (b= shape and steepness of germination curve, T_50_= time to 50% germination D_lag-50_= difference between time at germination onset (lag) and 50% germination, TMGR= time to maximum germination rate, Uniformity and AUC= area under curve) in unaged and aged seedlots at p≤0.05.

### Principal component analysis (PCA)

PCA was performed to examine the number of most informative principal components for the seven variables that accounts for most variability in the data for seventeen onion varieties. The three principal components (PC1 and PC2) taken together comprised 74.0% of the variability in the original data. PCA biplot defined PC1 (47.8%) and PC2 (26.2%) which altogether explained 74.0% of total variation for seven variables in the seventeen varieties (Fig. 4). The variables namely *TMGR*, *D*_*lag-50*_ and *T*_*50*_ were the major contributors to PC1, and for the PC2 *AUC* and *a* were the major players. The resulting PCA biplot revealed the correlation of the cluster analysis of seedlots into three different classes under distinct seed aging treatments by using the parameters extracted from 4-PHF.

**Fig 4:**
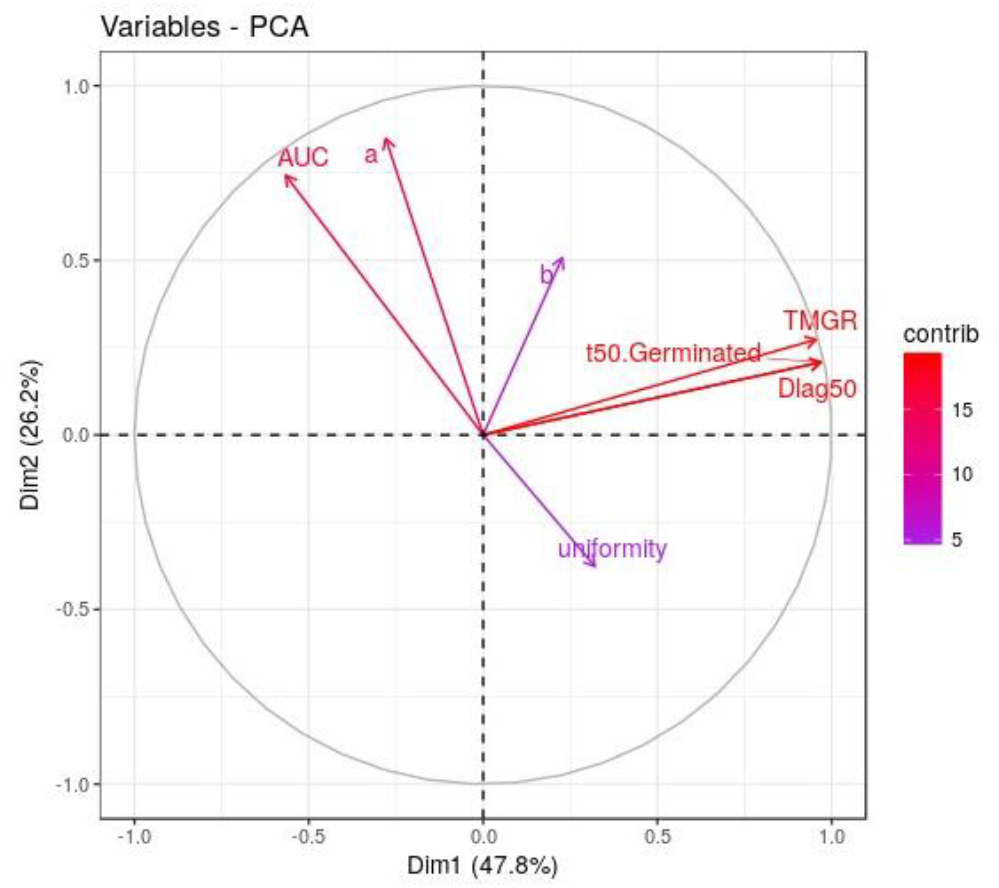
Biplot of seven parameters (*a, b*, *T*_*50*_, *D*_*lag-50*_, *TMGR, Uniformity and AUC*) extracted from 4-PHF in differentially aged 17 onion seedlots). The contribution to PC 1 is shown on the x-axis, while the contribution to PC 2 is on the y-axis. The values between the brackets indicate the percentage of the variance explained by the individual PCs. Each parameter contribution to the selected principle component is indicated by the length and color of the arrow.

### Cluster analysis

Based on the germination performance parameters extracted from the 4-PHF in unaged seedlots (fresh) (Fig 5: A.I & B. I) the following clusters were classified:

i. Cluster 1 (Unaged)-It comprised of Bhima Kiran, Bhima Shakti, Bhima Super, Bhima Shwetha, Bhima Shubhra, Arka Bheem and Arka Ujjwal varieties (Fig 5: A. I&B. I). The mean values of 4-PHF parameters showed that all varieties had the highest *a* (97±0.71%), *AUC* (1746.99±15.62), whereas the early germination parameters like *T*_*50*_ (1.88±0.05), *D*_*lag-50*_ (1.85±0.04), *TMGR* (1.58±0.02), and *U* (2.34±0.22) were lowest compared to other clusters (Table 2). This cluster was designated as “good storer” as all the seedlots possessed high viability/vigor
ii. Cluster 2 (Unaged)-Pusa Red, Pusa Madhavi, Pusa Riddhi, Pusa Early Grano, Agrifound Light Red, NHRDF Red and Arka Niketan were placed under Cluster 2 (Fig 5: A. I&B. I). Comparatively medium values of *a* (94±0.83), *AUC* (1446.92±12.75), were calculated for these varieties and it lay in between cluster 1 and 3. The time related parameters *T*_*50*_(4.68±0.14), *D*_*lag-50*_(4.49±0.14), *TMGR* (4.51±0.15) and *U* (2.17±0.16)showed that cluster 2 varieties required higher time to achieve the germination, therefore it was classified as “medium storers”(Table 2).
iii. Cluster 3 (Unaged)-Agrifound Dark Red, Arka Pragati and Arka Kalyan varieties were allocated to Cluster 3(Fig 5: A. I&B. I). It showed significantly lower *a* (92±2.38), *AUC* (1468.06±30.29), The time related parameters *T*_*50*_ (3.76±0.28), *D*_*lag-50*_ (3.59±0.32), *TMGR* (2.99±0.33) and *U* (4.97±0.28) led to its classification as “poor storers” (Table 2).

**Fig 5:**
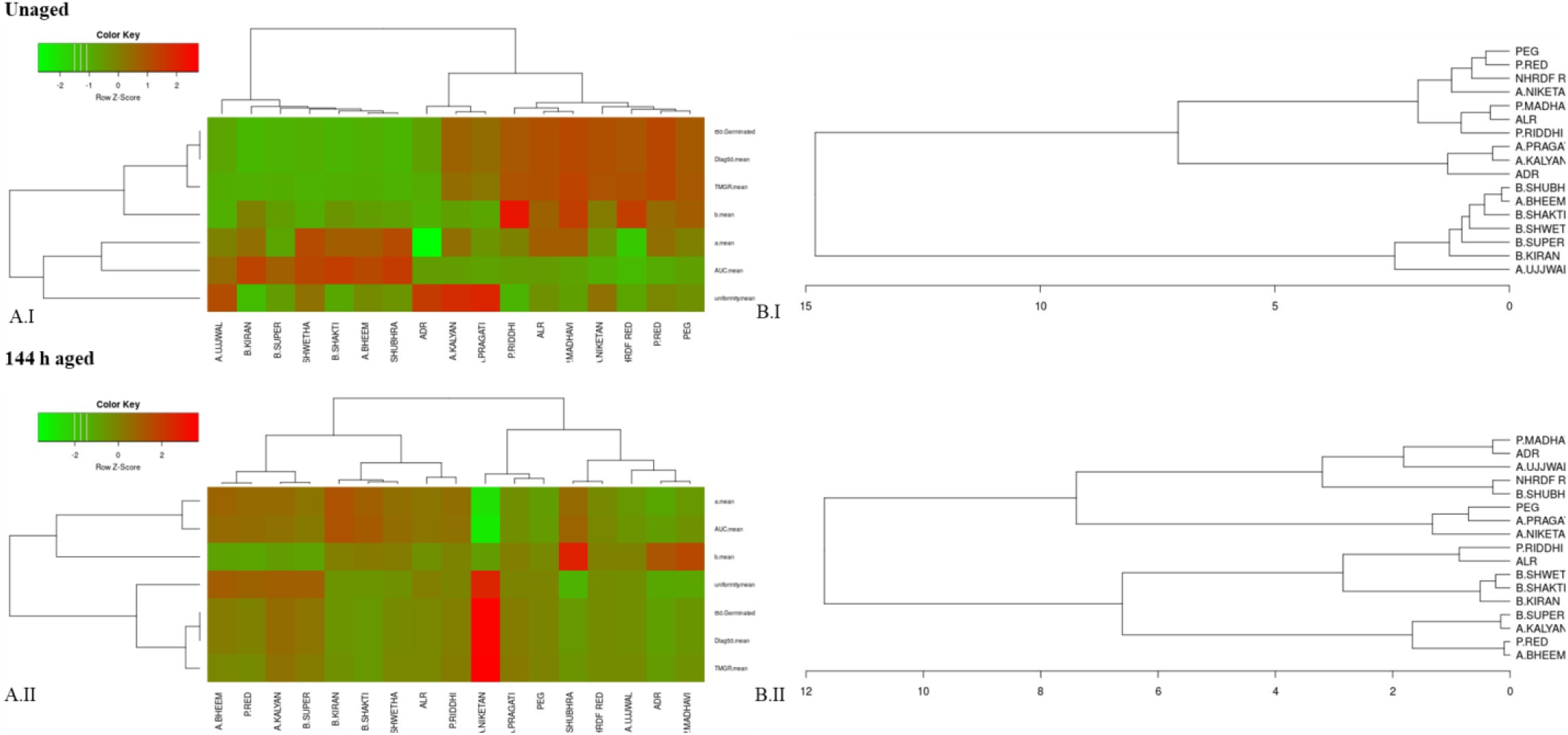
A. Heatmap (A) and dendrogram (B) generated from 4-PHF (*a, b*, *T*_*50*_, *D*_*lag-50*_, *TMGR, U, AUC*) for clustering of varieties of (I) Unaged (II) 144 h aged seedlots. The data was subsetted per treatment with columns representing the value of individual samples and rows selected parameter (*a, b*, *T*_*50*_, *D*_*lag-50, TMGR, U, AUC*_). Red and green represent high and low parameter value, respectively. The values of individual samples are normalized per parameter using z-Fisher transformation.

**Table2:**
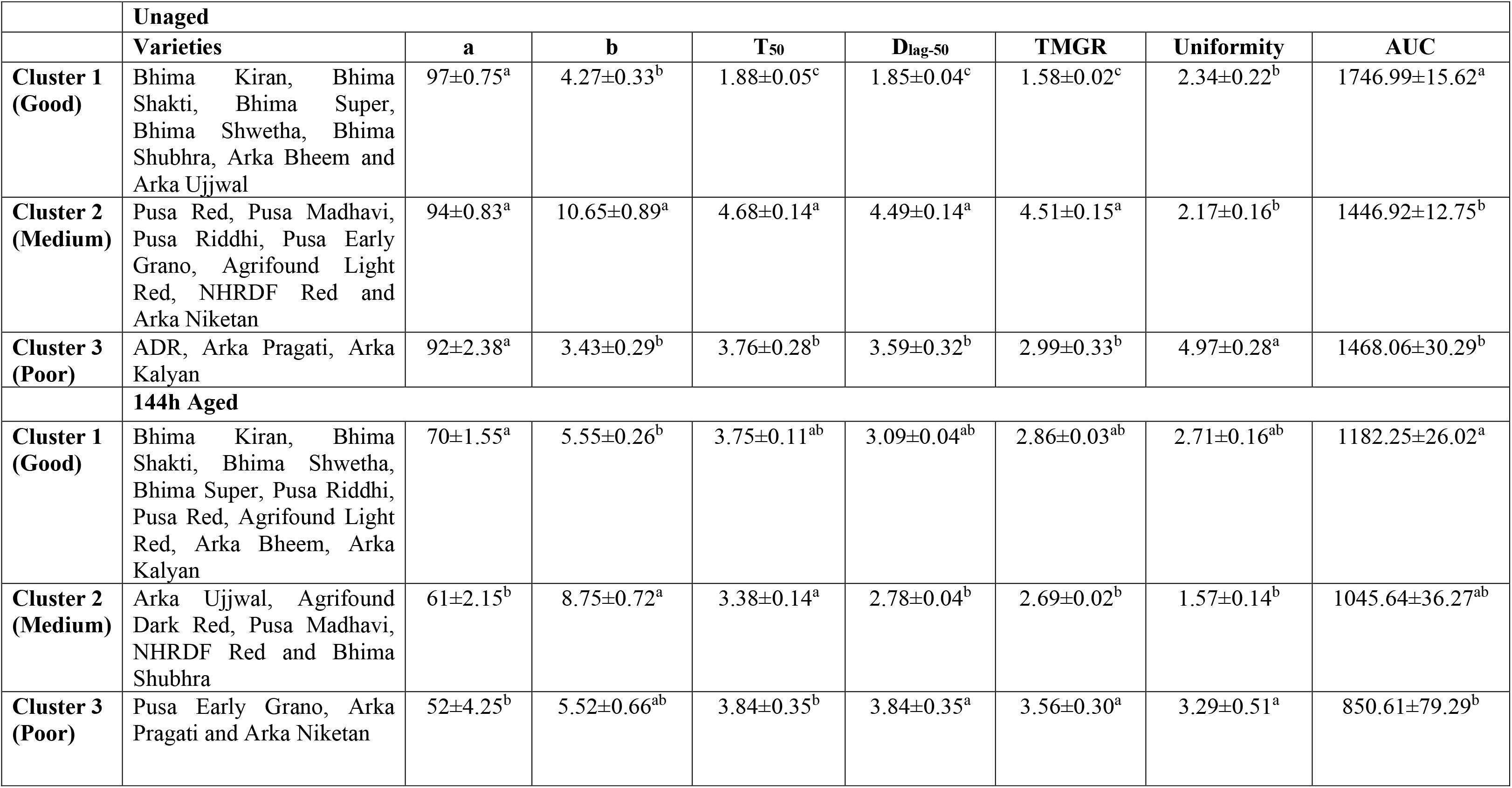
Mean values of different variables extracted from 4-PHF analysis used for classification of 17 seedlots of onion in different storability clusters (n=3). Cluster validation was performed by Tukey’s HSD pairwise test with p-value ≤ 0.05. Letters in superscript indicate significantly different groups

Upon 144h of ageing, clustering identified the following seedlots with differential storability (Fig 5: A. II&B. II)

i. Cluster 1(144 h aged)-It consist of Bhima Kiran, Bhima Shakti, Bhima Shwetha, Bhima Super, Pusa Riddhi, Pusa Red, Agrifound Light Red, Arka Bheem, Arka Kalyan (Fig 5: A.II&B. II). The mean values of 4-PHF parameters showed that all varieties had the highest *a* (70±1.55), *AUC* (1182.25±26.02), On the other hand, early germination parameters *T*_*50*_(3.75±0.11), *D*_*lag-50*_(3.09±0.04), TMGR (2.86±0.03),*U*(2.71±0.16)were lowest compared to other clusters (Table. 2). This cluster was designated as “good storer” as all the seedlots possessed high viability/vigor.
ii. Cluster 2-It consist of Arka Ujjwal, Agrifound Dark Red, Pusa Madhavi, NHRDF Red and Bhima Shubhra (Fig 5: A.II&B. II).Comparatively medium values of *a* (61±2.15%), *AUC* (1045.64±36.27), were calculated for these varieties and it lay in between cluster 1 and 3. Early germination parameters *T*_*50*_(3.75±0.11),and *D*_*lag-50*_(2.78±0.04), *TMGR* (2.69±0.02), *U*(1.57±0.14)(Table. 2). Although the time required to achieve 50% germination was less, the germination count was low, hence varieties were classified as “medium storer”.
iii. Cluster 3-It included Pusa Early Grano, Arka Pragati and Arka Niketan (Fig 5: A.II&B. II). It showed significantly lower *a* (52±4.25), *AUC* (850.61±79.29), Early germination parameters *T*_*50*_(3.84±0.35)and *D*_*lag-50*_(5.52±0.66), *TMGR* (3.56±0.30), *U*(3.29±0.51) required higher time to achieve 50% germination, therefore classified as “poor storer”(Table. 2).

Among seventeen seedlots, varieties from three seedlots namely Arka Niketan (poor storer), Pusa Madhavi (medium storer) and Bhima Kiran (good storer) were selected and the germination count data was fitted into 4-PHF curve. The corresponding area under curve (*AUC*) was a result of a graphical depiction of germination parameters (*a, b*, *T*_*50*_, *D*_*lag-50*_, *TMGR, U*) which allowed clear assessment of the relative storability group with and without ageing (Fig 6).

**Fig 6:**
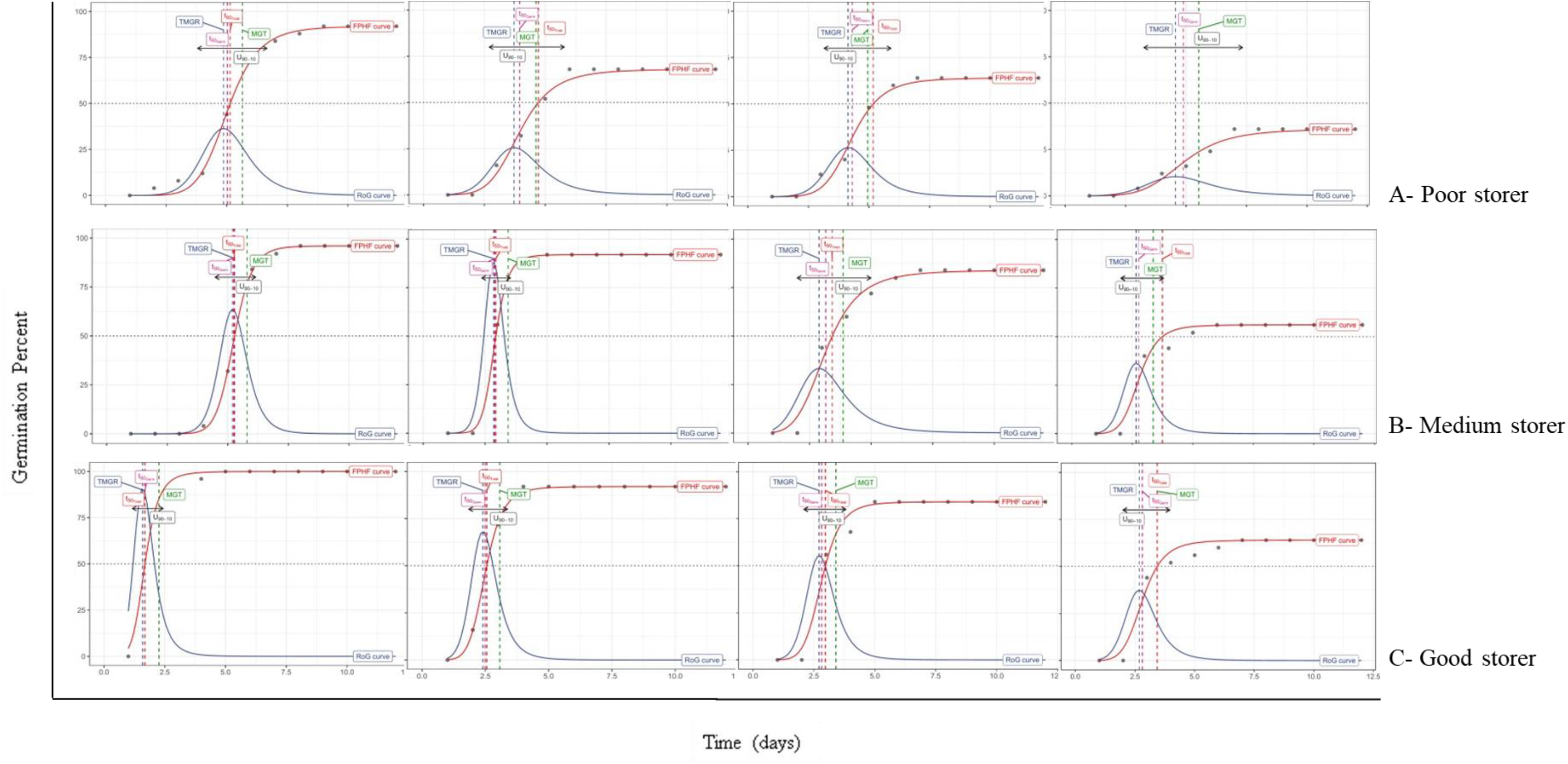
Curve fitting for germination count data of a representative sample of the identified good, medium and poor storers(A:Arka Niketan-Poor storer, B: Pusa Madhavi-Medium storer, C: Bhima Kiran: Good storer) using ‘FourPHFfit’ function of ‘germinationmetrics’ package in R program.

### Correlation among 4 PHF parameters

#### Unaged

Non-significant correlation was observed between *a, b*, *T*_*50*_, *D*_*lag-50*_, *TMGR* and uniformity.But *a* and *AUC* showed positive correlation (r=0.55**). Parameter *b*showedpositive correlation with early emergence related parameters *T*_*50*_(r=0.66**),*D*_*lag-50*_(r=0.66**) and *TMGR*(r=0.73**). Among the time related parameters highest correlation was observed between *TMGR* and both *D*_*lag-50*_(r=0.99**) and *T*_*50*_(r=0.99**). A non significant correlation was also observed between *U* and all other parameters (Fig 7.A).

**Fig7:**
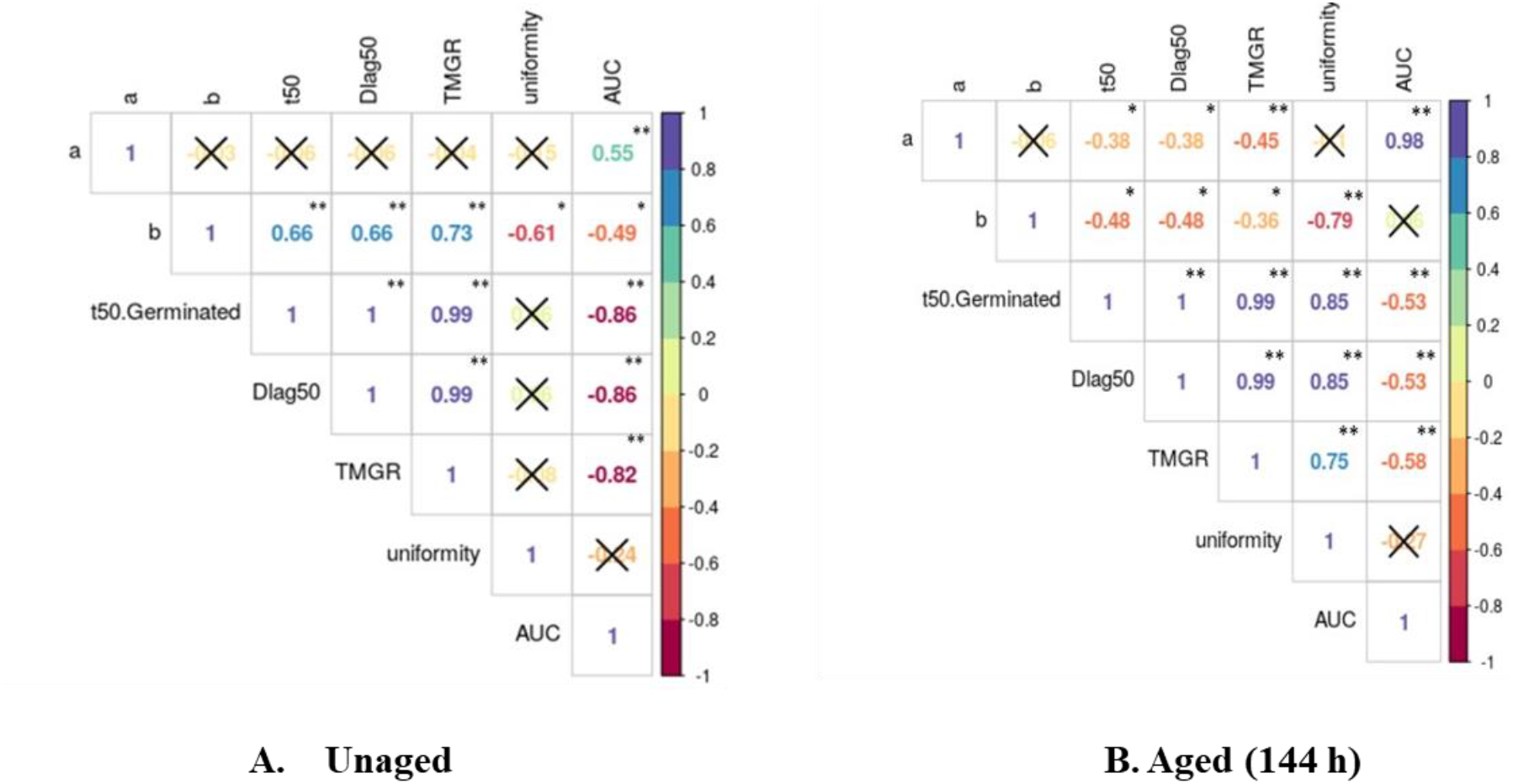
Pearson correlation coefficients between selected parameters under (A) Unaged (B) 144 h aged seeds. The color of the number reflects the strength of the correlation. The non-significant correlations, with the p-value above 0.05 are indicated with a X in the individual cells.

#### 144 h aged

Ageing treatment induced a homogeneity in correlation of various parameters. A high correlation (r=0.98**) existed between cumulative germination percent (*a*) and area under curve (*AUC*). The parameter *b*showed negative correlation with the other early emergence related parameters like*D*_*lag-50*_(r=−0.48) and *TMGR* (r=−0.36*). Anegative correlation was observed between the parameters*T*_*50*_*with* both *a* (r=−0.38), *AUC* (r=−0.53).Unlike unaged control, a significant correlation was also seen between *U* and *TMGR* (R=0.75**), *D*_*lag-50*_(r=0.85) and *T*_*50*_ (r=0.85)(Fig 7.B).

## Discussion

High seed quality is critical for deriving maximum profit from crop production. The establishment of successful crop stand determines the first step towards enhancing production and resource use efficiency. Seed vigour encompasses all the properties of the seed that are responsible for the performance of the seedlots of acceptable germination in a wide range of environmental conditions^22^. Correlating viability with vigor of seedlots has its limitations as seedlots with low viability can germinate faster and slow germinating ones can have high viability^23^. This gives a fallacious representation of the seedlot, the results of which are discernible only at harvest. Onion seeds are amongst the shortest-lived seeds that remain viable for eight to twelve months^24, 25^depending on the storage conditions. Storage of seedlots at high temperature and relative humidity, the conditions adopted in accelerated ageing test, lead to sequential loss of vigor and viability^26^. The present study was performed to classify 17 seedlots of Indian onion varieties into different storability groups (good, medium and poor storers) based on the germination related parameters. For this, the conventional approach of characterisation i.e. estimating germination capacity and vigor indices was compared against parameters extracted from 4-PHF. Significant differences between the various clusters dividing the varieties into good, medium and poor storers existed only for *SVI-I* values in the conventional approach. However, the extraction of germination count values under 4 PHF led to significant differences in germination values amongst the clusters demarcated in aged seedlots. This was due to difference in the set of varieties falling in the various clusters when the two approaches were compared. Classification using first approach revealed that varieties; Bhima Super and ADR that occupied Cluster 3 of “poor performers” under unaged conditions, shifted to cluster 1 of “good storers” after ageing. The probable underlying cause for this shift could be the modified structure of testa of these two varieties, radical scavenging facilitated by efficient antioxidant mechanism^27^or DNA and protein mechanism^28^ that resulted in better performance under ageing. In addition to these varieties, Arka Kalyan, Pusa Riddhi and Arka Bheem maintained a consistent response under both conditions and the deteriorative processes brought about stable changes that led to their medium performance under both conditions.

While using the extraction of parameters by 4-PHF, the varieties were found to shuffle between good and medium performers; and medium and poor performers when compared with their respective clusters in the conventional analysis. In unaged conditions, 4-PHF analysis classified Bhima Kiran, Bhima Shakti, Bhima Super, Bhima Shwetha, Bhima Shubhra, Arka Bheem, Arka Ujjwal as ‘good performers” as they exhibited high germination capacity (*a*), *AUC* and low early germination parameters (*D*_*lag-50*_, *TMGR*) and uniformity values. Lower *D*_*lag-50*_ and *TMGR* values suggested that the seedlot was vigorous and took less time to reach 50% germination. Even after 144h of ageing, the good performers could achieve high germination capacity with most of the seedlings emerging in 2.71 days within a small-time window. Minimising the spread of seedling emergence over time as evidenced by low *D*_*lag-50*_, *TMGR* values can help in establishment of uniform crop stand ensuring that most plants in the population attain similar size to contribute to the yield. Although time related parameters like *T*_*50*_, *D*_*lag-50*_, *TMGR* and uniformity were lower for medium performers (Cluster 2) than good ones under ageing, an approximate 11% decline in seed viability was observed. In order to avoid misinterpretation, the calculation of area under curve (*AUC*) for both the clusters was found to give the most comprehensive perspective of the performance of the seedlots as it takes into consideration all the extracted 4-PHF germination related parameters.

The germination count data of three clusters from 4-PHF was subjected to curve fitting to observe the trend of all the extracted parameters (representative samples from each cluster are illustrated in Fig 6). The ‘good performer’ showed high germination percentage with maximum seeds germinating within a short period of time. The progress of ageing led to faster flattening of the curve in poor performers, whereas good ones showed decrease in height of peak for rate of germination without its spreading. *TMGR* was also increased in ‘poor performers but the change was not remarkable in good ones. All these parameters clearly depicted that less area was covered under the curve in poor *vs* good storers.

Prognosis of performance under ageing by two approaches led to distribution of varieties in different clusters. Shifting between cluster 1---2, 2---3 and vice versa may not be of much significance but movement of varieties Pusa Red and Agrifound light red from poor performance cluster (Cluster 3 in conventional) to good performance (Cluster 1 of 4-PHF) in aged seed lots draws attention. Medium storers responded comparatively lower than good ones showing lower *a, AUC,* and longer *D*_*lag-50*_ and *TMGR* after 144 h of ageing. It was also noteworthy that Bhima Super occupied the ‘poor performer’ category under unaged condition but shifted to ‘good performer’ cluster under ageing in conventional classification, whereasin4 PHF classification it was good performer in both unaged and aged conditions. This change in its performance would have been unrecognised but for *a, TMGR* and *AUC* values of 68%, 2.99 days, 1113.5 respectively for parameters extracted from Hill function. All these values of Bhima Super were not significantly different from Cluster 1 (data not shown). The significance of *AUC* in determining the grouping of varieties to different clusters was reinforced with the change in germination characteristics of Bhima Super under ageing. Therefore, it could be concluded that final germination capacity lacks the ability to compare seedlots for storability as it highlights only the germination capacity and not the spread or speed of germination^17^. To avoid any misrepresentation of results, Kader (2005)^17^concluded germination index to be an inclusive parameter combining percentage of germination with speed of germination.

Irrespective of the seedlots, ageing time period showed significant difference at 96 and 144 h of ageing. The area under the curve which is a collective representation of parameters emanating by curve fitting from time zero to the time of final count was significantly different from control after 96 h of ageing.

In case of classification based on germination percentage and seedling vigor indices, interpretation oversees the crucial cumulative germination data over the germination period as a large number of samples are required to be analysed at different time intervals. Also, it does not take into consideration the details on start of germination and the rate of germination which are essential for a normally distributed population. Cultural practices like application of fertilizers, field maturity, umbel order and season of harvest are affected by the differences that arise due to duration of time between first and last seed to germinate, time needed for maximum number of seeds to germinate and difference in speed of germination^18^.Accelerated ageing shifted the varieties; Bhima Shubhra and Arka Ujjwal to the category of medium storers indicating that poor storage conditions can adversely affect the seeds by increasing lipid peroxidation, hampering seed coat structure and Amadori and Maillard reactions^29^. High temperature and humidity can damage DNA, RNA, protein and lipid after accumulation of ROS in seeds of these varieties^30, 31, 32^. This corroborates the previous report that suggest evaluation of germination performance after ageing to be the best criteria for determining relative storability of seedlots^33^. The varieties that maintain their status under unaged and aged conditions probably possess mechanisms to overcome the deterioration process which needs to be further investigated.

Correlation between the various parameters under unaged and aged conditions showed that *AUC* had a stronger negative correlation with time related parameters like *T*_*50*_, *TMGR* and *D*_*lag-50*_(r= −0.86,−0.82 and −0.86 respectively) in unaged whereas 144 h aged seeds were highly correlated with the germination percent (r=0.98**).This suggested that with the advent of deterioration processes, viability decreased (low *a* values) resulting in less *AUC* and longer time to accomplish 50% germination. The significantly positive correlation of *b* with *D*_*lag-50*_ and *T*_*50*_ in unaged and negative in aged seedlots respectively, indicated that the onset and speed of germination represented by the steepness and shape of curve and determined the time taken by the viable seeds to reach 50 % germination. It did not have any association with the germination performance of the unaged and aged seeds^21^. Uniformity did not show a significant correlation with any of the parameters under unaged condition but was significantly correlated to all parameters except germination count and AUC under aged condition. Ageing resulted in harmonizing the values in all the viable seeds of the seedlot based on their relative storability. Poor storers showed a non-uniform emergence i.e. spread of germination and seedling establishment as compared to good storers.

Thus, our study showed that the parameters extracted from 4-PHF were amenable to accurately defining the performance of seedlots under storage as mimicked by accelerated ageing conditions. It also highlighted the significance of using the parameter of area under curve (*AUC*) from the curve fitting graphs to allocate certain varieties (Bhima Super, Pusa Red and ALR in our study) to the appropriate storability class which would have otherwise been erroneously classified. The 4-PHF parameter could classify and predict storability of onion seed lots and could be successfully explored for other poor storer crops. Its usage can be extended for the studying storage behaviour of germplasm collections and plant genetic resources. Our study is the first reported research output of using 4-PHF mathematical function for seedlot characterization into different storability groups.

## Material and method

### Seed material

Freshly harvested seeds (2018-19) of seventeen onion varieties produced at ICAR-Indian Agricultural Research Institute, New Delhi (Pusa Riddhi, Pusa Red, Pusa Madhavi, Pusa Early Grano), ICAR-Indian Institute of Horticultural Research, Bengaluru (Arka Pragati, Arka Kalyan, Arka Bheem, Arka Ujjwal, Arka Niketan), ICAR-Directorate of Onion and Garlic Research, Pune (Bhima Kiran, Bhima Shakti, Bhima Shwetha, Bhima Shubhra, Bhima Super) and National Horticultural Research and Development Foundation, New Delhi (NHRDF Red, Agrifound Dark Red, Agrifound Light Red) were used for the study.

### Seed conditioning and accelerated ageing

The initial moisture content of fresh onion seedlots ranged from 4-7% which were brought to 13-15% by conditioning seeds for three days with saturated solution of sodium chloride (40mg/100ml) in desiccators (76% R. H.). Seedlots were artificially aged by maintaining temperature (42°C) and relative humidity (100%) throughout the ageing period^22^ for 144h and samples were drawn at 48h interval i.e., at 0 h, 48h, 96h and 144h. After ageing treatment, the seeds were dried back to their initial moisture content.

### Standard germination test

Eight replicates of fifty seeds were planted on moist blotter paper in a Petridish placed in a germinator maintained at 20 °C temperature and 90% relative humidity^22^. Germination count was recorded up to 12 days. The germination count data was fitted to curve using 4-PHF.

### Components of 4-Parameter Hill Function (4-PHF)

The cumulative germination count data of onion seedlots from 0 (first count) to 12 days (final count) was fitted to 4-PHFusing the “germination metrics package” from R programme. The following equation was used for calculating 4-PHF^21^,

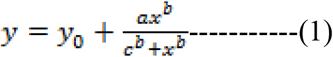

Where, the variables y and x are cumulative germination percent and standard germination test duration respectively, y_0_ is intercept on the y-axis.

After fitting the germination count data to the above equation 1, the FourPHFfit extracts the major germination parameters. Different functions were collectively used in the FourPHFfit to compute and extract the following parameters (2 to 6)^21,34^.

1. **Germination percent/capacity (*a*)** Maximum germination achieved by a seed lot which is represented as cumulative germination percent in the curve was determined.
2. **Shape or steepness of germination curve (*b*)** The *b* value represents the shape or steepness of germination curve.
3. **Time to 50% germination (*T*_*50*_)** It was calculated as the time required to achieve 50% of seed germination in the total seeds.
4. **Difference between time at germination onset (*lag*) and 50% germination (*D*_*lag-50*_)** The time difference between the onset of germination (lag) and time to reach 50% germination (*T*_*50*_) was calculated under this parameter as shown in equation (2). It indicates uniformity, earliness of germination and vigor. Smaller the D_lag-50_ value earlier the germination, uniformity and higher steepness of germination curve.

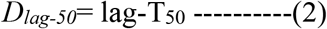
5. **Time to maximum germination rate (*TMGR*)** Daily rate of germination was considered instead of the mean germination rate. Maximum germination rate at a specific point of time is called as instantaneous rate of germination. It was computed by partial derivation of equation (1)

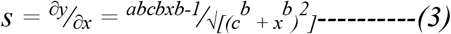

*s* in the equation (3) represents daily rate of germination. After plotting a graph between the *s* and time, point of time at which the maximum germination rate is reached is defined as TMGR shown in equation (4). It’s another early germination parameter which means smaller the TMGR value shorter the time to reach T_50_, more vigorous is the seed lot.

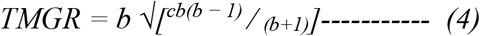
6. **Uniformity (*U*)** It’s the time required to achieve the percent of germination in argument.
7. **Area under curve (*AUC*)** The measurement of *AUC* considers the germination capacity (a), uniformity, early germination parameters like *TMGR*, *T*_*50*_, *D*_*lag-50*_ and *b* (shape and steepness of germination curve) collectively. It was estimated by integrating the curve fitting from time zero (0) to the time of final count (argument t_max_).

### Seed vigour indices

Ten normal seedlings were selected randomly after standard germination test. The germination percent, seedling length (cm) and dry weight (mg) were recorded. The seed vigour indices were assessed as a product of germination (%) and seedling length (cm)/ dry weight (mg) and calculated by using the formula given by Abdul-Baki and Anderson, 1973^35^.

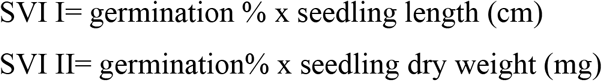

### Data analysis

Germinationmetrics package available in R program was used to visualize the data and extract the parameters from germination count data^34^. The ANOVA test, complemented by a pairwise Tukey’s HSD test was used to examine significant differences among the different treatments. MVApp^36^, a multivariate analysis application was used to conduct principal component and cluster analysis for classifying seventeen onion seedlots into different categories based on relative storability.

## Acknowledgements

The Senior Research Fellowship provided by ICAR-Indian Agricultural Research Institute, New Delhi, India to the first author during his doctoral studies is acknowledged.

## Author information

### Author notes

**Sunil Kumar**

**Present address:** PhD Scholar, Division of Seed Science and Technology, ICAR- Indian Agricultural Research Institute, Pusa Campus, New Delhi 110012, India

**Sudipta Basu**

**Present address:** Principal Scientist, Division of Seed Science and Technology, ICAR- Indian Agricultural Research Institute, Pusa Campus, New Delhi 110012, India

**Anjali Anand**

**Present address:** Principal Scientist, Division of Plant Physiology, ICAR- Indian Agricultural Research Institute, Pusa Campus, New Delhi 110012, India

**Aravind J.**

**Present address:** Scientist, Division of Germplasm Conservation, ICAR- National Beaureu of Plant Genetic Resources, Pusa Campus, New Delhi 110012, India

### Affiliations

Division of Seed Science and Technology, Division of Plant Physiology, ICAR-Indian Agricultural Research Institute, New Delhi, 110 012, India

Division of Germplasm Conservation, ICAR- National Bureau of Plant Genetic Resources, Pusa Campus, New Delhi 110012, India

Sunil Kumar, Sudipta Basu, Anjali Anand, J. Aravind

### Contributions

SB and AA conceptualized the hypothesis, designed the experiment and finalized the manuscript.SK performed the experiment, analysed the data and drafted the manuscript. SK and JA carried out data analysis and visualization in R. All authors reviewed the manuscript.

## Ethics declarations

### Competing Interests

The authors declare no competing interests.

## Notes

### Competing Interest Statement

The authors have declared no competing interest.

## References

1. Walters, C. Orthodoxy, recalcitrance and in-between: describing variation in seed storage characteristics using threshold responses to water loss. Planta 242, 397–406(2015).

2. Rajjou, L. & Debeaujon, I. Seed longevity: survival and maintenance of high germination ability of dry seeds. C. R. Biol. 331(10), 796–805. 10.1016/j.crvi.2008.07.021 (2008).

3. Walters, C. Understanding the mechanisms and kinetics of seed ageing. Seed Sci. Res.8, 223 – 244, https://doi.org/10.1017/S096025850000413X(1998).

4. Groot, S., Surki, A.A., Vos, R.& Kodde, J. Seed storage at elevated partial pressure of oxygen, a fast method for analysing seed ageing under dry conditions. Ann. Bot.110, 1149 – 1159, https://doi.org/10.1093/aob/mcs198 (2012).

5. Hourston, J. E. et al. The effects of high oxygen partial pressure on vegetable Allium seeds with a short shelf-life. Planta 251 (6),105 https://doi.org/10.1007/s00425-020-03398-y (2020).

6. Bentsink, L. et al. Genetic analysis of seed-soluble oligosaccharides in relation to seed storability of Arabidopsis. Plant Physiol. 124 (4),1595–1604, https://doi.org/10.1104/pp.124.4.1595 (2000).

7. Job, C., Rajjou, L., Lovigny, Y., Belghazi, M., & Job, D. Patterns of protein oxidation in Arabidopsis seeds and during germination. Plant Physiol. 138 (2),790–802, https://doi.org/10.1104/pp.105.062778(2005).

8. FAOSTAT. Primary crops http://faostat.fao.org/faostat/form?collection = Production.Crops. Primary & Domain = Production & servlet = 1&hasbulk = 0&version = ext & language = EN (2017).

9. MoA & FW. Ministry of Agriculture and Farmers Welfare, Government of India https://http://agricoop.nic.in/sites/default/files/Horticulture%20Statistics%20at%20a%20Glance-2018.pdf (2018).

10. Mohamed-Yasseen, Y., Costanza, S., & Splittstoesser, W. Onion seed anatomy in relation to aging. J Veg. Crop Prod.1(2),51–69, https://doi.org/10.1300/J068v01n02_05(1996).

11. Amalfitano, C.et al. Yield, antioxidant components, oil content, and composition of onion seeds are influenced by planting time and density. Plants 8(8),293, https://doi.org/10.3390/plants8080293 (2019).

12. Sano, N. et al. Staying alive: molecular aspects of seed longevity. Plant Cell Physiol. 57, 660–674(2016).

13. Ranal, M. A. & Santana, D. G. D. How and why to measure the germination process. Braz. J. Bot.29(1),1–11(2006).

14. Labouriau, L. G. A germinação das sementes. Série de Biologia. Monografia, 24 (1983).

15. Brown, R. F. & Mayer, D. G. Representing cumulative germination. 1. A critical analysis of single-value germination indices. Ann. Bot. 61(2),117–125(1988a).

16. Bewley, J.D. & Black, M. Seeds: Physiology of development and germination. (2nd ed. Plenum Press, New York) (1994).

17. Kader, M. A. A comparison of seed germination calculation formulae and the associated interpretation of resulting data. J. and Proc. the Royal Society of New South Wales138, 65–75(2005).

18. Kader, M. & Jutzi, S. Drought, heat and combined stresses and the associated germination of two sorghum varieties osmotically primed with NaCl. Phytogen3, 22–24 (2001).

19. Tipton, J.L. Evaluation of three growth curve models for germination data analysis. J. Amer. Hort. Soc.109, 451–454 (1984).

20. Brown, R.F. & Mayer, D.G. Representing cumulative germination. 2. The use of the Weibull function and other empirically derived curves. Ann. Bot.61, 127–138, https://doi.org/10.1093/oxfordjournals.aob.a087535 (1988b)

21. El-Kassaby, Y. A., Moss, I., Kolotelo, D. & Stoehr, M. Seed germination: mathematical representation and parameters extraction. For. Sci.54(2),220–227(2008).

22. ISTA. International rules for seed testing. The International Seed Testing Association (ISTA), Baaserdorf, Switzerland (2015).

23. Thomson, A.J., & Y.A. El-Kassaby. Interpretation of seed-germination parameters. New For.7, 123–132 (1993).

24. Thirusendura Selvi, D. & Saraswathy, S. Seed viability, seed deterioration and seed quality improvements in stored onion seeds: a review J. Hort. Sci. Biot.93(1),1–7, https://doi.org/10.1080/14620316.2017.1343103(2018).

25. Roberts, S. Vegetable seed quality, storage and handling: AHDB Factsheet. Agriculture and Horticulture Development Board, Kenilworth, vol 3(2018).

26. Tian, X., Song, S.& Lei, Y. Cell death and reactive oxygen species metabolism during accelerated ageing of soybean axes. Russ. J. Plant Physiol.55 (1),33–40 (2008).

27. Bailly, C., Benamar, A., Corbineau, F. & Côme, D. Free radical scavenging as affected by accelerated ageing and subsequent priming in sunflower seeds. Physio. Plant.104(4),646–652(1998).

28. Fu, Y. B., Ahmed, Z. & Diederichsen, A. Towards a better monitoring of seed ageing under ex situ seed conservation. Con. Physiol. 3(1),10.1093/conphys/cov026 (2015).

29. Narayana Murthy, U. M., Prakash P. K. & Sun, W. Q. Mechanisms of seed ageing under different storage conditions for *Vigna radiata* (L.) Wilczek: lipid peroxidation, sugar hydrolysis, Maillard reactions and their relationship to glass state transition. J. Expt. Bot.54(384),1057–1067, https://doi.org/10.1093/jxb/erg092 (2003).

30. Harman, D. “Free radical theory of aging: history,” in Free Radicals and Aging, (eds I. Emerit and B. Chance) 1–10. (Basel: Birkhäuser-Verlag, 1992). https://doi.org/10.1007/978-3-0348-7460-1_1

31. Neelesh Kapoor, Arvind Arya, Mohd. Asif Siddiqui, Hirdesh Kumar and Asad Amir. Physiological and biochemical changes during seed deterioration in aged seeds of rice (*Oryza sativa* L.). Amer. J. Plant Physiol.6, 28–35, https://doi.org/10.3923/ajpp.2011.28.35(2011).

32. Ratajczak, E., Małecka, A., Bagniewska-Zadworna, A., and Kalemba, E. M. The production, localization and spreading of reactive oxygen species contributes to the low vitality of long-term stored common beech (*Fagus sylvatica* L.) seeds. J. Plant Physiol.174, 147–156. doi: 10.1016/j.jplph.2014.08.021 (2015).

33. Delouche, J. C. & Baskin, C. C. Accelerated aging techniques for predicting the relative storability of seedlots. Seed Sci. & Technol.1, 427–452(1973).

34. Aravind, J., Vimala, D., Radharani, J., Jacob, S. R. & Srinivasa, K. “The germinationmetrics package: A brief introduction.” New Delhi, India: ICAR-National Bureau of Plant Genetic Resources.https://github.com/aravindj/germinationmetricshttps://cran.rproject.org/package=germinationmetrics(2019).

35. Abdul-Baki, A.A. & Anderson J.D. Vigor determination in soybean seed by multiple criteria. Crop Sci.13, 630–633, https://doi.org/10.2135/cropsci1973.0011183X001300060013x(1973).

36. Julkowska, M. M. et al. MVapp—multivariate analysis application for streamlined data analysis and curation. Plant Physiol. 180(3),1261–1276, https://doi.org/10.1104/pp.19.00235(2019).

